# Salivary and plasma levels of matrix metalloproteinase-9 and myeloperoxidase at rest and after acute physical exercise in patients with coronary artery disease

**DOI:** 10.1101/455915

**Authors:** Zeid Mahmood, Helena Enocsson, Maria Bäck, Anna K Lundberg, Lena Jonasson

## Abstract

**Background:** Low-grade systemic inflammation is a predictor of recurrent cardiac events in patients with coronary artery disease (CAD). Plasma proteins such as matrix metalloproteinase (MMP)-9 and myeloperoxidase (MPO) have been shown to reflect basal as well as stress-induced inflammation in CAD. Measurements of MMP-9 and MPO in saliva might pose several advantages. Therefore, we investigated whether salivary levels of MMP-9 and MPO corresponded to plasma levels in patients with CAD, both at rest and after acute physical exercise.

**Methods:** An acute bout of physical exercise on a bicycle ergometer was used as a model for stress-induced inflammation. Twenty-three CAD patients performed the test on two occasions 3-6 months apart. Whole unstimulated saliva was collected before, directly after and 30 min after exercise while plasma was collected before and after 30 min. MMP-9 and MPO in saliva and plasma were determined by Luminex.

**Results:** MMP-9 and MPO levels were 2- to 4-fold higher in saliva than in plasma. Within the saliva compartment, and also to a great extent within the plasma compartments, MMP-9 and MPO showed strong intercorrelations at all time points. However, there were no (or weak) correlations between salivary and plasma MMP-9 and none between salivary and plasma MPO.

**Conclusion:** We conclude that salivary diagnostics cannot be used to assess systemic levels of MMP-9 and MPO in CAD patients, neither at rest nor after acute physical exercise.

## Introduction

Inflammation is an important component of atherosclerosis, from the formation of atherosclerotic plaques to plaque destabilization eventually leading to plaque rupture and atherothrombotic events, such as myocardial infarction [1]. Systemic levels of inflammatory markers can be useful to assess cardiovascular risk and monitor disease activity and over the years, great efforts have been made to identify relevant and easily available markers. The predictive value of markers like C-reactive protein (CRP) and interleukin (IL)-6 in determining the risk of myocardial infarction is well-documented [2]. Moreover, a number of epidemiological and clinical studies have shown that neutrophil-associated proteins in plasma, such as matrix metalloproteinase (MMP)-9 and myeloperoxidase (MPO), predict cardiovascular outcome [3, 4] and also relate to the extent and severity of atherosclerosis [5]. Overall, there is emerging evidence that the presence of low-grade systemic inflammation should be considered in clinical praxis when assessing a patient’s risk of recurrent cardiovascular events.

Low-grade inflammation is however not always detectable in plasma measures taken at rest. As has been shown in various settings, stress provocation tests may add important information on the individual’s susceptibility to inflammatory response [6-9]. A greater inflammatory response to daily stressors has also been proposed to render atherosclerotic plaques unstable and more prone to rupture [10]. Measurements of MMP-9 and MPO may be useful in the assessment of stress-induced inflammatory response since they are stored in neutrophils and rapidly released upon stressful stimuli. E.g., neutrophils from patients with coronary artery disease (CAD) are more prone to release these mediators, in particular MMP-9, upon in vitro stimulation compared with neutrophils from healthy subjects [11]. Moreover, patients with CAD who exhibit a significant and early increase in plasma MMP-9 after a laboratory stress test show signs of more advanced disease, including premature cellular aging and larger atherosclerotic burden [12].

As an alternative or complement to blood-based tests, salivary diagnostics has emerged as a promising tool in assessing inflammation. Compared with blood, saliva provides some distinct advantages such as non-invasiveness, ease and multiple sampling opportunities. So far, much focus has been on saliva as a diagnostic tool for oral disease but growing evidence indicates that it can be useful also for systemic diseases [13-15]. Interestingly, a few studies have indicated that salivary levels of MMP-9 and MPO are potentially promising biomarkers in cardiovascular disease. When Floriano et al [16] used a saliva-based nano-biochip test for the diagnosis of acute myocardial infarction, they found that both MMP-9 and MPO were significantly elevated in saliva collected within 48 h of chest pain onset. Another study demonstrated a significant association between salivary levels of MMP-9 and subclinical cardiovascular disease, more precisely carotid intima-media thickness, in 250 individuals [17]. However, none of these studies included plasma or serum measurements and it has remained largely unclear whether salivary levels of MMP-9 and MPO reflect systemic levels.

The aim of the present study was to investigate whether salivary levels of MMP-9 and MPO in patients with CAD corresponded to systemic levels. We measured the concentrations in saliva and plasma, under basal conditions and after an acute bout of physical exercise, on two separate occasions 3-6 months apart.

## Methods

### Study population

Twenty-three patients with a recent coronary event, i.e. acute coronary syndrome and/or revascularization with either percutaneous coronary intervention or coronary artery bypass grafting, were consecutively recruited from the Outpatients’ Cardiology Clinic at the University Hospital in Linköping, Sweden. Exclusion criteria were age > 75years, severe heart failure, neoplastic disease, major clinical depression, chronic liver and renal failure, chronic immunologic disorders or treatment with immunosuppressive/anti-inflammatory agents, serious physical or psychological disease interfering with performing an exercise test, and inability to understand the Swedish language. The study was conducted in accordance with the ethical guidelines of Declaration of Helsinki, and the research protocols were approved by the Ethical Review Board in Linköping. Written informed consent was obtained from all patients and controls.

### Exercise stress test

The patients underwent exercise stress tests on 2 occasions; 3-4 weeks (visit A) and 4-6 months (visit B) after the index coronary event. A standardized submaximal cycle ergometer test was used according to the WHO protocol [18], with an increased workload of 25 W every 4.5 min. The initial starting load, 25 W or 50 W, was decided based on the patient’s exertion history. A pedalling rate of 50 rates per minute was used. After 2 and 4 minutes at each work load, heart rate, rating of perceived exertion according to Borg’s rating of perceived exertion scale (RPE) [19] and subjective symptoms, including chest pain and dyspnea according to Borg’s Category Ratio Scale, CR-10, scale were rated. After 3 minutes, the systolic blood pressure was registered. The exercise test was discontinued at Borg RPE 17 and/or dyspnea 7 on Borg’s CR-10 scale. This submaximal exercise test on a bicycle ergometer has excellent test-retest reliability in patients with CAD [20] and is a well-established, safe procedure that is frequently used in exercise-based cardiac rehabilitation in Sweden.

### Sampling procedure

The patients first lay down on a bed reclining comfortably for 20-30 min. Thereafter, saliva samples were collected prior to the bicycle test, directly after and 30 min after the completion of the test whereas blood samples were collected prior to and 30 min after the test. Saliva samples for basal measurements were always taken before initiating venepuncture to avoid a possible stress effect. Patients had been asked to avoid tooth brushing, smoking, eating and drinking for at least 2 h before sample collection. The saliva was collected with Salimetrics Oral cotton Swabs (Salimetrics via Electrabox, Stockholm, Sweden) placed under the tongue for 2 minutes. The saliva samples were then immediately placed on ice. Whole blood was collected in 9 mL EDTA tubes (BD biosciences) and plasma was obtained after centrifugation for 10 min at 1500 x g. Saliva and plasma samples were frozen at -70 °C.

### Measurement of MMP-9 and MPO in plasma

MMP-9 and MPO in plasma was measured by magnetic bead-based luminex assay (R&D Systems, Minneapolis, USA) following the manufacturers’ instructions. All samples were assayed in a dilution of 1:50. The interassay coefficients of variation (CV) were 13.8 % for MMP-9 and 15 % for MPO. The limits of detection were 0.136 ng/mL for MMP-9 and 0.148 ng/mL for MPO.

Since plasma volume (PV) loss can be a confounding variable in the interpretation of changes in plasma concentration, the values after exercise were corrected for PV loss. Hematocrit values before and after exercise were assayed from whole blood using a standard cell counter. PV values were defined as (1 – hematocrit) and thereafter percent change in PV or delta PV (ΔPV) from rest to exercise was calculated. To adjust for PV loss, MMP-9 and MPO values after exercise were multiplied by (1 +(ΔPV/100)) [21].

### Measurement of MMP-9 and MPO in saliva

MMP-9 and MPO in saliva were measured by the same kit as described above, i.e. magnetic bead-based luminex assay (R&D Systems). Briefly, saliva samples were gently thawed in 4°C and thereafter centrifuged 10 000 x g for 5 minutes. All samples were assayed in a dilution of 1:50 and were re-run at 1:100 or 1:10 if outside the standard curve. For all patients except one, basal and post-exercise saliva were assayed at the same plate. To correct for differences in saliva volume (e.g. due to dry mouth after the exercise test), the concentrations of MMP-9 and MPO were normalized to total protein content in saliva using the Bradford protein assay (Bio-Rad, Solna, Sweden). All samples were run in duplicate, and diluted 1:5 or 1:3 prior to the Bradford assay. If samples were outside the standard curve, they were re-run in 1:10 dilution or undiluted. The interassay CV was 8.5 % for MMP-9 and 7.2 % for MPO.

### Statistics

SPSS 25 for Windows (IBM Corp. Armonk, NY) was used for statistical analyses. For clinical and laboratory characteristics, data are presented as median (25th-75th percentiles). The statistical significance of any difference between two groups was tested by using Mann-Whitney U-test. Chi-square test was used for nominal data. Wilcoxon signed-ranks test was used for pair-wise comparisons and Friedman test for more than two correlated samples. Spearman’s rank correlation coefficient was used for correlation analyses. A p-value < 0.05 was considered statistically significant.

## Results

### Basal characteristics and exercise testing

The basal characteristics of the 23 patients, including cardiovascular medication and laboratory data at visit A, are summarized in Table 1. The cardiovascular response during the exercise test at visit A is also shown in Table 1. The heart rate and systolic blood pressures increased significantly during cycling (p < 0.001). At 30 min after exercise, the systolic blood pressure returned to basal levels while the heart rate was still significantly higher compared to basal heart rate (p < 0.001). The duration of exercise was 15 (12-18) min and a maximal watt level of 125 (100-125) was achieved. Among the 23 patients who participated in visit A, 19 participated in visit B while 4 declined to participate. The cardiovascular response, duration of exercise and exertion level obtained at visit B were similar to those at visit A (data not shown).

**Table 1.** Basal characteristics of patients, including demographic and clinical data, cardiovascular medication, laboratory data and cardiovascular response during acute physical exercise at visit A.

|  |  |  |
| --- | --- | --- |
| Age, years |  | 65 (60-69) |
| Males, n (%) |  | 18 (75) |
| Smokers, n (%) |  | 5 (21) |
| Body mass index, kg/m<sup>2</sup> |  | 25.9 (23.4-27.9) |
| *Index event* |  |  |
| Acute coronary syndrome<sup>a</sup>, n (%) |  | 15 (65) |
| Stable angina, n (%) |  | 8 (35) |
| *Medication* |  |  |
| Dual anti-platelet therapy<sup>b</sup>, n (%) |  | 22 (92) |
| Beta-blocker, n (%) |  | 18 (75) |
| ACE-I/ ARB<sup>c</sup>, n (%) |  | 18 (75) |
| Calcium-channel blocker, n (%) |  | 1 (4) |
| Statins, n (%) |  | 24 (100) |
| *Laboratory data* |  |  |
| Total cholesterol, mmol/L |  | 3.5 (3.1-4.3) |
| C-reactive protein, mg/L |  | 1.0 (0.4-1.7) |
| *Cardiovascular response to acute physical exercise* |  |  |
| Heart rate,<br>beats/min | Basal | 63 (55-72) |
|  | Peak | 121 (111-131)** |
|  | 30 min recovery | 69 (60-83)** |
| Systolic blood pressure,<br>mm Hg | Basal | 130 (115-140) |
|  | Peak | 193 (175-203)** |
|  | 30 min recovery | 120 (115-130) |
| Maximal level of perceived exertion (Borg RPE)<sup>d</sup> |  | 17 (16-17) |

Values are given as median (25<sup>th</sup> – 75<sup>th</sup> percentiles) or n (%). \*\* p < 0.001 compared to basal levels, <sup>a</sup>non-ST-elevation myocardial infarction or ST-elevation myocardial infarction, <sup>b</sup> low-dose aspirin + clopidogrel or ticagrelor, <sup>c</sup> angiotensin converting enzyme inhibitors/angiotensin receptor blockers, <sup>d</sup> Borg's rating of perceived exertion scale, from 6 to 20 where 17 means "very hard exertion".

### Salivary levels of MMP-9 and MPO

The total protein content in saliva and protein-adjusted salivary levels of MMP-9 and MPO before (i.e. basal) and after exercise stress tests on two test occasions (visit A and B) are presented in Table 2. The total protein levels in saliva increased significantly directly after exercise and returned to basal levels 30 min after exercise, probably reflecting a reduced salivary flow during exercise. The protein-adjusted levels of MMP-9 and MPO in saliva did not differ significantly between rest and exercise, neither did levels differ between visit A and B. However, without adjusting for total protein content, the salivary levels of MMP-9 and MPO showed significant increases after exercise as shown in Figure 1 a and b. In the following text, the salivary levels of MMP-9 and MPO will be referred to as protein-adjusted levels.

**Table 2.** Measurement in saliva and plasma before (basal), directly after (peak) and 30 min after (post) acute physical exercise.

|  |  |  | Visit A | Visit B | p<sup>a</sup> |
| --- | --- | --- | --- | --- | --- |
| **Total protein** | Saliva,<br>*μg/mL* | Basal | 754 (410-1421)<sup>x</sup> | 728 (573-1365)<sup>x</sup> | 0.586 |
|  |  | Peak | 1508 (946-2325) | 1284 (890-1785) | 0.286 |
|  |  | Post | 852 (598-1220)<sup>x</sup> | 881 (653-1168)<sup>x</sup> | 0.872 |
|  |  | p<sup>b</sup> | <0.001 | <0.001 |  |
| **MMP-9** | Saliva<br>*pg/mg total protein* | Basal | 92 (53-211) | 104 (51-295) | 0.492 |
|  |  | Peak | 103 (52-229) | 104 (42-342) | 0.647 |
|  |  | Post | 141 (84-279) | 117 (82-271) | 0.936 |
|  |  | p<sup>b</sup> | 0.260 | 0.327 |  |
|  | Plasma<br>*pg/mL* | Basal | 50 (37-82) | 43 (28-84) | 0.601 |
|  |  | Post | 50 (25-73) | 51 (22-91) | 0.334 |
|  |  | P<sup>a</sup> | 0.287 | 0.904 |  |
| **MPO** | Saliva<br>*pg/mg total protein* | Basal | 82 (31-222) | 124 (32-378) | 0.084 |
|  |  | Peak | 118 (47-317) | 99 (422-375) | 0.557 |
|  |  | Post | 109 (42-203) | 107 (43-256) | <u>0.573</u> |
|  |  | p<sup>b</sup> | 0.084 | 0.662 |  |
|  | Plasma<br>*pg/mL* | Basal | 27 (22-36) | 23 (17-29) | <u>0.036</u> |
|  |  | Post | 27 (21-35) | 24 (18-28) | <u>0.227</u> |
|  |  | p<sup>a</sup> | 0.638 | 0.673 |  |

Salivary levels of total saliva and protein-adjusted levels of MMP-9 and MPO were measured before (basal), directly after an acute bout of exercise (peak) and 30 min after exercise (post). Plasma levels of MMP-9 and MPO were measured before and 30 min after exercise, the latter adjusted for plasma loss. Exercise tests were performed on two occasions (visit A and B). Values are given as median (inter-quartile range). <sup>a</sup> Wilcoxon signed-ranks test were used for comparisons between visit A and B, <sup>b</sup> Friedman's test for comparisons between basal, peak exercise and post-exercise levels, <sup>c</sup>and Wilcoxon signed-ranks test for comparisons between basal and post-exercise levels.

**Figure 1a and b.**
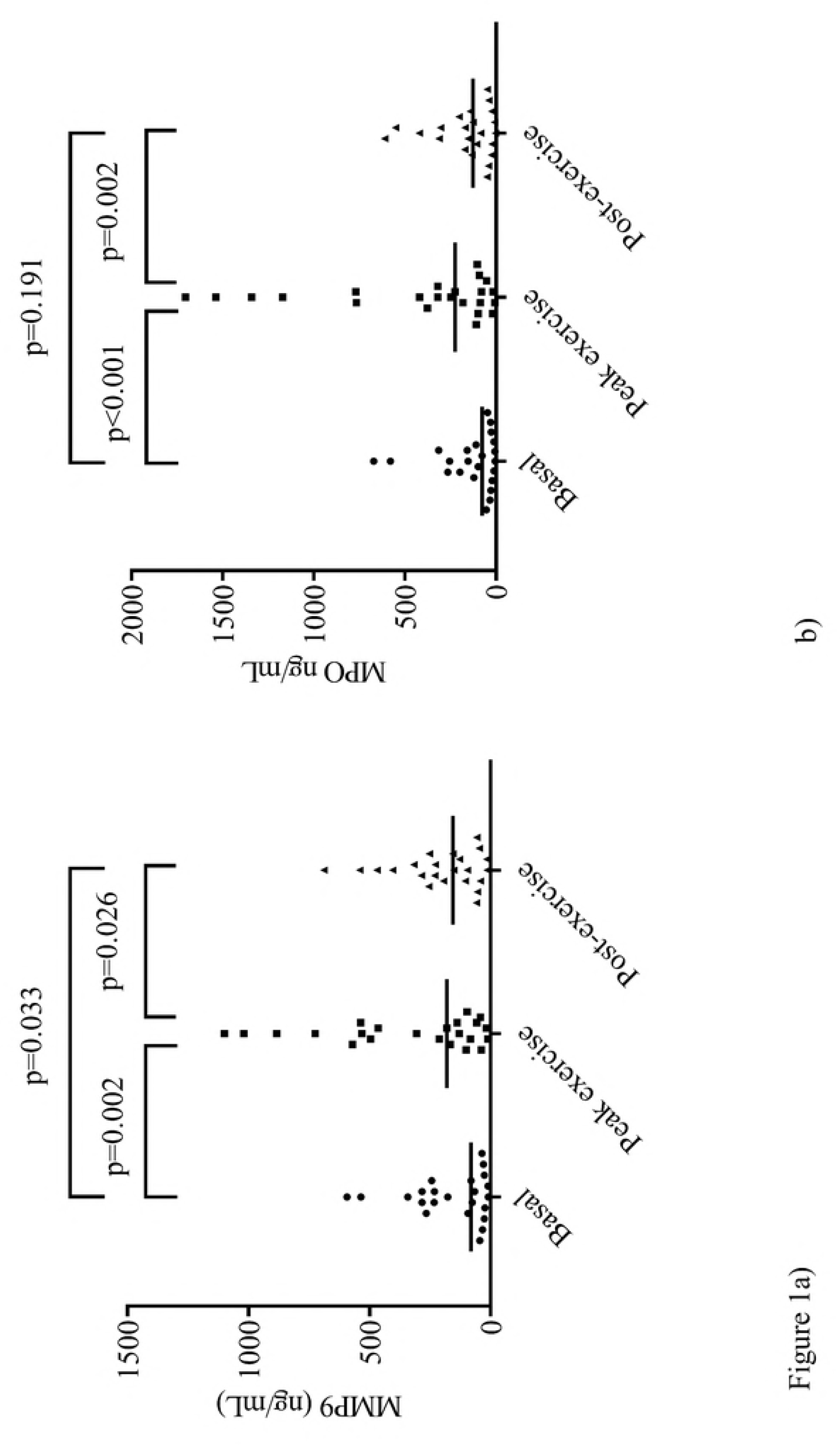
Crude values of MMP-9 and MPO before and after an exercise stress test. MMP-9 (1a) and MPO (1b) in saliva are presented as crude values, i.e. without correction for total protein content, before (basal), directly after exercise (peak exercise) and 30 min after exercise (post-exercise). Horizontal lines represent median values.

### Plasma levels of MMP-9 and MPO

The levels of MMP-9 and MPO were in general lower in plasma than in saliva, around 2-fold and 4- fold, respectively. Plasma levels of MMP-9 and MPO before and after acute exercise (the latter corrected for PV loss) are presented in Table 2. There were no significant differences, neither between rest and exercise, nor between visit A and B. The ΔPV values were relatively low on both test occasions, visit A -1.6 (-1.8-1.7) % and visit B -1.7 (-3.3-0.00). Hence, uncorrected levels of MMP-9 and MPO after exercise did not differ significantly from PV-corrected levels (data not shown).

### Correlations between MMP-9 and MPO in saliva and plasma

The correlations between MMP-9 and MPO in saliva and plasma before and after exercise on the two test occasions are presented in Table 3 and 4, respectively. Within the saliva compartment, the levels of MMP-9 were highly intercorrelated. The levels of MMP-9 were also correlated within the plasma compartment, though to a lesser extent than in saliva. There were either no or weak correlations between MMP-9 levels in plasma and saliva (Table 3). The levels of MPO were highly correlated both within the saliva and plasma compartments whereas no correlations were seen between salivary and plasma levels (Table 4). Further, within the saliva compartment, the MMP-9 and MPO levels were highly intercorrelated, visit A; basal *r* = 0.707, peak *r* = 0.693, and post-exercise *r* = 0.750, respectively, all p < 0.001. Also, within the plasma compartment, the MMP-9 and MPO levels correlated with each other at visit A; basal, *r* = 0.461, p = 0.027 and post-exercise, *r* = 0.593, p = 0.003, respectively. Similar correlations between MMP-9 and MPO within the saliva and plasma compartments were seen at visit B (data not shown).

**Table 3.** Correlations of MMP-9 levels in saliva and plasma before(basal) and 30 min after acute physical exercise (post).

|  | Saliva A<br>basal | Saliva A<br>post | Saliva B<br>basal | Saliva B<br>post | Plasma A<br>basal | Plasma A<br>post | Plasma B<br>basal | Plasma B<br>post |
| --- | --- | --- | --- | --- | --- | --- | --- | --- |
| Saliva A<br>basal | 1 | 0.751*** | 0.807*** | 0.719** | 0.157 | 0.286 | 0.332 | 0.088 |
| Saliva A<br>posts |  | 1 | 0.899*** | 0.908*** | 0.252 | 0.416 | 0.414 | 0.243 |
| Saliva B<br>basal |  |  | 1 | 0.847*** | 0.077 | 0.497* | 0.584* | 0.377 |
| Saliva B<br>post |  |  |  | 1 | 0.202 | 0.434 | 0.533* | 0.317 |
| Plasma A<br>basal |  |  |  |  | 1 | 0.380 | 0.368 | 0.620** |
| Plasma A<br>posts |  |  |  |  |  | 1 | 0.633** | 0.624** |
| Plasma B<br>basal |  |  |  |  |  |  | 1 | 0.785*** |
| Plasma B<br>post |  |  |  |  |  |  |  | 1 |

The table shows correlations (Spearman's rho) between MMP-9 in saliva (adjusted for total protein) and plasma before (basal) and 30 min after acute physical exercise (post), the latter adjusted for plasma loss, on two test occasions (A and B). \* $p < 0.05$ , \*\* $p < 0.01$ , \*\*\* $p <$ 0.01.

**Table 4.** Correlations of MPO levels in saliva and plasma before (basal) and 30 min after acute physical exercise (post).

|  | Saliva A<br>basal | Saliva A<br>post | Saliva B<br>basal | Saliva B<br>post | Plasma A<br>basal | Plasma A<br>post | Plasma B<br>basal | Plasma B<br>post |
| --- | --- | --- | --- | --- | --- | --- | --- | --- |
| Saliva A<br>basal | 1 | 0.685*** | 0.841*** | 0.807*** | 0.010 | 0.102 | 0.297 | 0.226 |
| Saliva A<br>Post |  | 1 | 0.605** | 0.635** | 0.260 | 0.352 | 0.376 | 0.346 |
| Saliva B<br>basal |  |  | 1 | 0.836*** | 0.061 | 0.221 | 0.432 | 0.169 |
| Saliva B<br>post |  |  |  | 1 | 0.147 | 0.294 | 0.383 | 0.463 |
| Plasma A<br>basal |  |  |  |  | 1 | 0.945*** | 0.767*** | 0.841*** |
| Plasma A<br>post |  |  |  |  |  | 1 | 0.886*** | 0.856*** |
| Plasma B<br>basal |  |  |  |  |  |  | 1 | 0.895*** |
| Plasma B<br>post |  |  |  |  |  |  |  | 1 |

The table shows correlations (Spearman's rho) between MPO in saliva (adjusted for total protein) and plasma before (basal) and 30 min after acute physical exercise (post), the latter adjusted for plasma loss, on two test occasions (A and B). \*\* p < 0.01, \*\*\* p < 0.01.

In line with previous results from our group (12), the stress-induced release of MMP-9 in plasma showed a large variation among patients with CAD. We therefore divided the patients into two groups depending on stress-induced release of plasma MMP-9. At visit A, 7 patients (30 %) exhibited an increase in plasma MMP-9 after exercise, +70 (16-106) %, while the remaining 16 patients showed a decrease, -31 (-45-(-15)) %. This was however not reflected in the saliva. The two groups showed similar and non-significant changes in salivary MMP-9 after exercise, 31 (-19-55) % and 47 (-26-92) %, respectively, p = 0.974, without any correlation to changes in plasma.

## Discussion

During the last decades, substantial improvements have been achieved in medical as well as lifestyle management of CAD, all of which may lead to reductions in inflammatory activity. Despite this, a residual inflammatory risk for recurrent cardiac events exists in many patients with CAD [22]. There is thus an urgent need to identify this risk population. However, inflammatory markers in plasma and serum samples that are collected during resting conditions do not always provide the full picture. The inflammatory state may not become evident until the patient is exposed to stress. Both psychosocial and exercise-related stressors can be used to elicit an inflammatory response. Comparisons between CAD patients and healthy controls have shown that exposure to any of these stressor types results in larger CRP increases in patients than in controls [6, 7]. However, there are some methodological concerns when using CRP in stress tests. Due to the response time of hepatic CRP release, it may take between 6 and 24 h before any changes in CRP appear. In contrast, neutrophil-associated proteins like MMP-9 and MPO are rapidly released in response to stress. Recently, we were able to show that a significant increase in MMP-9 30 min after a psychological stress test occurred in one third of patients with CAD [12]. Similarly, 30 % of the patients in the present study showed a significant increase in MMP-9 30 min after an exercise stress test. So far, we can only speculate that the stress-induced level of MMP-9 is a marker of residual inflammatory risk. However, the association between stress-induced release of MMP-9 and shorter leukocyte telomere length as well as increased plaque burden support the hypothesis that recurring bouts of inflammation in response to exogenous challenges contribute to a gradual progression in CAD [12].

Saliva has emerged as a non-invasive tool for assessing MMP-9 and MPO in cardiovascular disease [16, 17, 23]. If salivary levels reflect systemic levels, this might open up for a new and simple way to monitor inflammation, in particular stress-induced inflammation, in patients with CAD. Therefore, the main aim of this study was to investigate the relationships between salivary and plasma concentrations of MMP-9 and MPO in patients with CAD. This was performed on two separate occasions 3-6 months apart, at rest as well as after an exercise stress test. In saliva, the proteins were highly intercorrelated with their respective counterparts at all time points, and, to a great extent, they were also intercorrelated in plasma. However, there were no or a few weak correlations between the analytes in saliva and plasma. This argues for a distinct oral compartment in which MMP-9 and MPO reside independently from blood. Only a few previous studies have compared salivary and systemic levels of MMP-9 and MPO, though with disparate results. Foley *et al* [23] performed serial measurements of MMP-9 and MPO in serum and unstimulated whole saliva after alcohol septal ablation in 21 patients with obstructive hypertrophic cardiomyopathy. They reported significant correlations between serum and salivary levels of MMP-9 and MPO at all time points. Another study by Meschiari *et al* [24], including 23 patients with periodontal disease and 19 healthy controls, found significant correlations between plasma levels of MMP-9 and gelatinolytic activity in saliva samples collected by expectoration after chewing on paraffin (i.e. stimulated whole saliva) [24]. Two possible reasons for the discrepancy between the findings in our study and previous studies deserve particular mentions. First, serum levels of neutrophil proteins may not be representative of systemic levels due to the degranulation of neutrophils and platelets during the ex vivo blood clotting process in the collection tubes. Secondly, none of the previous studies presented protein-adjusted salivary levels. The salivary concentrations of proteins can vary in response to stimulation and alteration of saliva flow. In the present study, we observed higher concentrations in saliva directly after exercise if we did not adjust for total protein.

The high correlations between salivary levels of MMP-9 and MPO and their respective counterparts on two different test occasions suggest that salivary levels are rather stable over time. It has been pointed out that the day-to-day fluctuation of inflammatory markers in saliva may be substantial due to intraindividual variability. However, the variation is greater with at-home collection compared with collection in the clinic under supervision [25]. In a previous study, Fingleton *et al* [26] reported that the within-subject variability in salivary MMP-9 activity was relatively small when 5 samples were collected over a 3-week period.

The present study is the first to measure the degree to which salivary MMP-9 and MPO levels correspond to plasma levels in response to exercise-induced stress. However, a number of smaller studies have investigated the reliability of saliva-based to blood-based levels of cytokines, like IL-1β and IL-6, in response to acute stress (either social-cognitive or exercise-physical). As summarized in a review by Slavish *et al* [27], salivary levels of cytokines do not appear to be good approximates of what is happening systemically. Salivary and plasma levels of cytokines correlate poorly with each other before as well as after stress. One probable explanation is that cytokines are too large to enter saliva via passive diffusion. The high molecular weights of MMP-9 and MPO, 82 and 140 kDa, respectively, may thus partly explain why the salivary levels do not correlate with the corresponding plasma levels.

Our study has a number of potential limitations that need to be considered. First, we did not control for oral health status. Periodontitis is known to be overrepresented among patients with CAD [28]. Both MMP-9 and MPO are abundantly present in periodontal tissue and salivary levels have been shown to correlate with periodontal inflammation [24, 29]. Still, there is some controversy over whether salivary MMP-9 levels are higher in patients with periodontitis. In a recent study of patients with and without periodontitis, no significant differences between groups were found in either serum or salivary levels of MMP-9 [30]. Another limitation of our study is the small sample size not allowing us to evaluate the impact of potential confounders, such as smoking. Moreover, the present results may only apply to patients with CAD. The patients were taking a number of different medications, like statins and platelet inhibitors, that are associated with anti-inflammatory effects [31, 32].

## Conclusion

MMP-9 and MPO were abundantly present in saliva of CAD patients with concentrations exceeding those in plasma. However, there was no evidence that salivary levels reflected circulating levels, neither at rest nor after acute physical exercise. As regards MMP-9 and MPO, saliva should rather be seen as a separate compartment and not as a mirror of systemic disease. Salivary diagnostics does not seem to be a useful alternative to plasma diagnostics in the assessment of systemic levels of MMP-9 and MPO in CAD patients.

## Acknowledgements

This research was funded by the Heart Lung Foundation, Sweden (Grant number: 20150648) and Swedish Research Council, Sweden (Reference number: 2014-2479). The funding sources had no role in study design, data collection and analysis or preparation of the manuscript.

